# TreeTOP: Plant experimental platforms in canopy space

**DOI:** 10.64898/2026.08.30.748063

**Authors:** Julia Baumeister, Moe Bakhtiari, Mona Schreiber, Michael Eisenring, Martin M. Gossner, Susanne Walden, Anke Becker, Marie-Lara Bouffaud, Simone Cesarz, Benjamin Dauphin, Nico Eisenhauer, Kezia Goldmann, Lea Heidrich, Stephanie Jurburg, Robert Junker, Jürgen Kreuzwieser, Christian Lampei, Thomas Nauss, Martina Peter, Luis Prada-Salcedo, Mika Tarkka, Christiane Werner, Dirk Zeuss, Sylvie Herrmann, Francois Buscot, Katrin Heer, Lars Opgenoorth

## Abstract

1. Forest canopies harbour strong microclimatic gradients that shape plant performance, species interactions and ecosystem processes. Yet, despite renewed interest sparked by global change, forest canopies remain difficult-to-access experimental spaces.
2. With the goal to expand access to tree canopies as experimental arenas, we designed, built, and tested TreeTOP, a standardized experimental platform that opens canopy space for manipulative ecological experiments, specifically with potted plants. TreeTOP features lightweight aluminum frames placed in mature tree canopies non-invasively, allowing potted plants to be placed in three different heights, ground level, shade canopy, and sun canopy.
3. We implemented TreeTOP using two contrasting infrastructure concepts to demonstrate its applicability in both highly equipped canopy research facilities and forests without permanent canopy infrastructure. One installation relied on a canopy crane, grid power and fully automated irrigation, whereas the second was built by certified tree climbers and was equipped with an autonomous solar-powered, battery-operated irrigation system. At both sites, environmental sensor networks monitor the experiment.
4. TreeTOP successfully reproduced characteristic canopy microclimatic gradients, including increasing light availability, daytime air temperatures and thermal extremes with canopy height. Despite differing infrastructures, both implementations generated comparable microclimatic patterns, demonstrating that standardized canopy experiments are feasible in forests with or without permanent canopy access. By opening canopy space for manipulative experiments, TreeTOP provides a transferable framework for investigating plant performance, phenology, species interactions and microbiome assembly under realistic forest conditions.

## 1 INTRODUCTION

Forest canopies form the primary interface between forest ecosystem and the atmosphere and are therefore directly exposed to ongoing climate change (Ozanne et al. 2003, Nakamura et al. 2017). Widespread canopy dieback across diverse tree species following recent heat waves, droughts, and global-change related emerging native and non-native pests and pathogens have renewed interest in the investigation of canopy ecology (De Frenne 2024, Simler-Williamson et al. 2019, Verheyen et al. 2024, Nolan et al. 2021, Walthert et al. 2021, Nakamura et al. 2017, Sallé et al. 2021, Sire et al. 2022).

Canopies are not homogenous entities but harbor various gradients in environmental conditions related to insolation, wind speed, air temperature, vapor pressure deficit, leaf temperature, and gas exchange, which all act on different spatiotemporal scales (Vinod et al. 2023). These variable factors have direct impacts on known ecophysiological traits such as leaf economics spectrum traits, photoprotection, VOC emission, as well as gas exchange (Vinod et al. 2023). However, while many within-canopy trait gradients have been well described, we often lack mechanistic understanding of what drives specific traits, as various biophysical factors overlap and phenotypic traits interact. An additional challenge arises from the fact, that trees are vertically structured modular organisms whose canopies comprise assemblages of repeated modules, such as shoots bearing groups of leaves (Harper et al. 1986). In deciduous trees, new shoot-leaf modules are produced each year, in some species through multple flushes within a single growing season (Späth 1912). These modules develop within the described environmental heterogeneity of the crown and may thereofore acquire distinct morphological, physiological, and chemical properties depending on their position during development. This modularity creates a fundamental challenges for mechanistic canopy ecology: differences observed among naturally developed modules at different canopy positions may reflect their developmental history, their current environment, or both. Observational comparisons alone therefore cannot readily disentangle the effects of canopy position and local environmental conditions from phenotypic differences established during module development.

Even less is known about how these gradients individually and interactively shape the traits and community structure of canopy-inhabiting species. For example, it has recently been shown that trees harbor distinct tissue-specific and canopy position-specific microbiomes (Arnold et al. 2025). Likewise, insect community composition and performance have been shown to vary within forest canopies (Ulyshen 2011; Seifert et al. 2020; Eisenring et al. 2021). How these components collectively shape the interactions between trees and the organisms associated with them is far less well understood. For example, do solar-radiation-induced leaf physical properties such as thicker cuticles reduce herbivory pressure? Or are these patterns driven by nutrient availability, defensive phytochemicals, or abiotic conditions? Recent studies have provided first mechanistic insights into how within-canopy environmental heterogeneity shapes canopy-associated communities. For example, microbiomes from sun- and shade-exposed leaves have been shown to differ not only in composition but also in their functional effects on host performance (Saueressig et al. 2026).

Together, these studies demonstrate both the promise of functional canopy research and our still limited ability to disentangle the relative contributions of host traits, developmental history, microclimate variation, dispersal ability, and other biotic and abiotic drivers to community assembly under natural forest conditions. Addressing these questions requires manipulative approaches that complement observations of naturally developed canopy modules by exposing comparable plant material to different positions within mature tree canopies. Such approaches provide an experimental framework in which the initial state and subsequent developmental environment of plant material can be standardized or deliberately varied.

Here, we describe TreeTOP – a platform installed in tree crowns to place tree microcuttings or saplings into different canopy heights and thereby open canopy space for manipulative ecological experiments. We contrast a high-cost canopy-crane based approach with a low-cost flexible tree-climber based approach. As one of the main challenges in tree-canopy experiments is to supply water to the experimental plants, we also contrast an automatized electric-grid based irrigation system with a low-cost battery self-supported system. Also, we give insights into sensor systems for monitoring these setups and quantify costs and effort for the setup. We discuss the effectiveness of the platform to compare canopy differences and discuss common and infrastructure-specific pitfalls in the designs. Finally, we provide future directions on how these experimental studies can advance the mechanistic understanding of how climate change and its effects on within canopy/tree crown heterogeneity affect ecological interactions and processes.

## 2 Material and methods

TreeTop was set up in two experimental research forests. First, the Marburg Open Forest is a 244-ha research- and teaching-dedicated forest of the Philipps University Marburg in central Germany. Located within the subcontinental beech forest zone (212–488 m a.s.l.), it is dominated by European beech (*Fagus sylvatica*) with sessile oak (*Quercus petraea*), accounting for approximately 30% of the forest area. Over the past years, the Marburg Open Forest has been developed into a multidisciplinary forest research site hosting long-term climatological, hydrological, and tree physiological sensor networks, biodiversity monitoring infrastructure, and regular unmanned aerial vecile (UAV) sensing campaigns (Zeuss et al. 2023, www.uni-marburg.de/MOF). Second, the Research Forest in Hölstein is an experimental forest located in the Jura mountains of Switzerland (540 m a.s.l.) and is managed by the University of Basel. The 1.6-ha comprises a mature, species-rich temperate mixed forest with more than 400 trees representing 14 species. It hosts the Swiss Canopy Crane II, a 50-m-tall canopy crane providing access to approximately 250 trees from 12 species. The forest is equipped with extensive long-term research infrastructure including automated tree physiological, soil, and microclimatic sensor networks. Established in 2018 as a long-term climate change experimental incubator, the site is designed to investigate the impacts of increasing drought on mature temperate forests through large-scale rainfall exclusion and continuous ecosystem monitoring (https://ppe.duw.unibas.ch/en/sccii/).

### 2.1 Experimental platform construction

The individual platforms were designed to fit six 12 l plant pots each, with a pot diameter and height of 26 cm. For a lightweight construction, each platform was welded using 2.5 cm x 2.5 cm aluminum square tubes, resulting in a metal frame that was 90 cm long, 62 cm wide, and 28 cm high, with cross-bracing added for more stability and to hold the pots in place (figure 1). To later facilitate a straightforward placement of plants on the platform and to prevent any movement of the pots during stormy conditions, pots were chosen that have a cross-shaped recess that exactly fit the cross-bracing and holes for excess water. Empty pots were tightly connected to the platform. For this connection, a hole was screwed in the middle of the bottom of each empty pot, which was fitted on a bolt and secured with a metal plate and nut to safely secure it onto the platform. Exactly fitting inlet pots were used to fill with soil and place the plants into. These were then transferred into the empty pots. See supplementary material for specifications.

**Figure 1.**
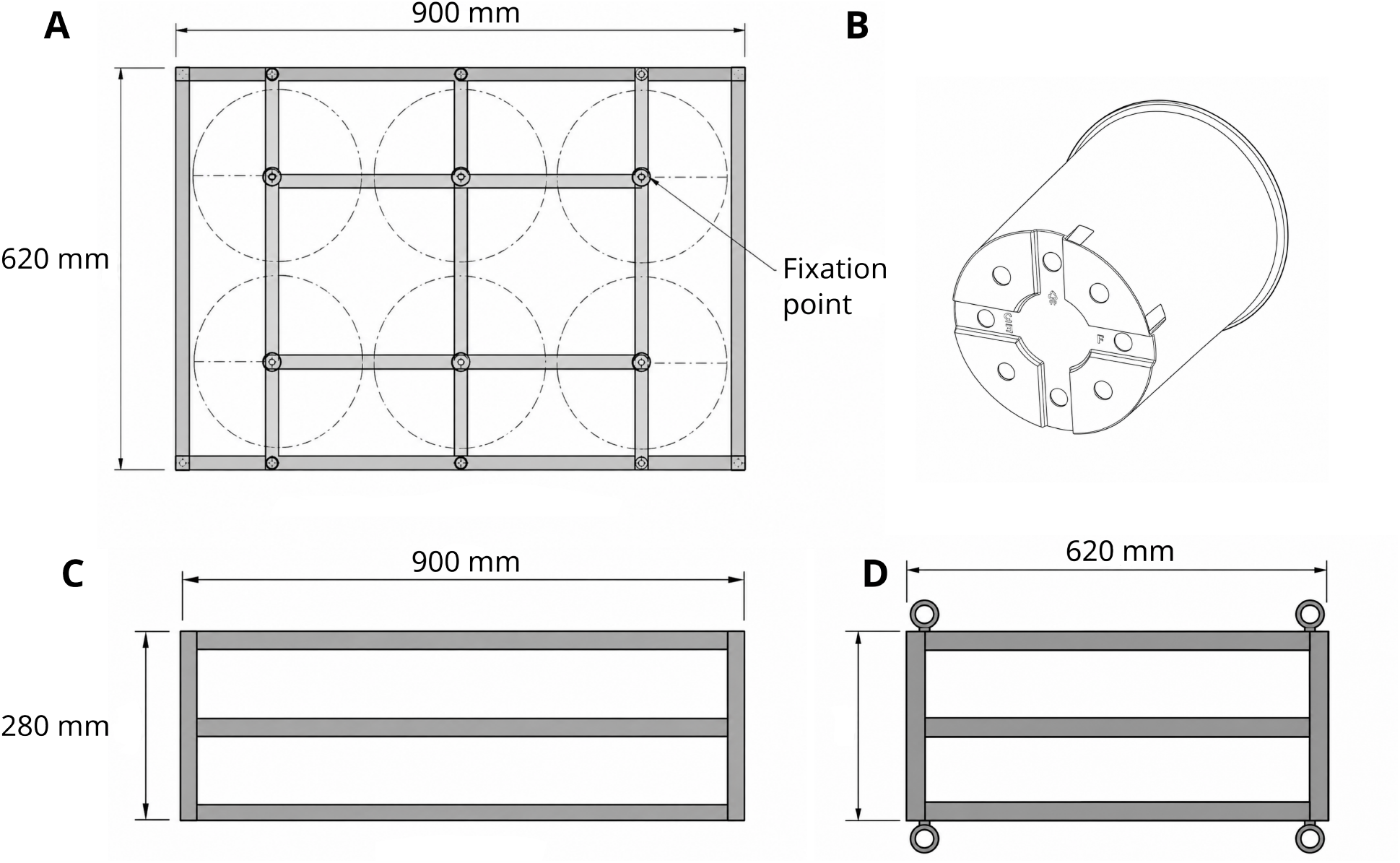
Experimental platform construction and pot attachment system. Technical drawings of the aluminum frames used to accommodate six 12-L plant pots. (A) Top view showing the 900 × 620 mm frame, cross-bracing, and arrangement of the six pots. (B) Bottom view of a plant pot showing the cross-shaped recess designed to fit onto the platform bracing. (C) Side view of the 900 × 280 mm platform. (D) Front view showing the 620 × 280 mm frame. Platforms were constructed from 2.5 × 2.5 cm aluminum square tubing. Pots were secured to the frame using a central bolt, metal plate, and nut.

### 2.2 Platform placement in the canopy

Experimental platforms were established at three vertical positions: ground level, middle canopy, upper canopy. At ground level, pots were placed adjacent to the respective experimental tree, whereas platforms in the middle and upper canopy were attached to the tree stem. For the placement of the experimental platforms in the upper and middle canopy heights, two different strategies were employed, either using a canopy crane (Swiss Canopy Crane II, Hölstein, Switzerland) or relying on tree climbers (Marburg Open Forest, Germany). The placement of platforms in the upper canopy was mainly determined by reachability by the crane or tree climbers. For the middle canopy, the section underneath the main tree crown was chosen as position for the platform. Table 1 shows an overview of each tree and the heights of experimental platfom placements.

**Table 1.** Heights of the TreeTOP platforms installed in the middle and upper canopy of the six experimental trees at the Hölstein Forest and the Marburg Open Forest. Vertical distance refers to the difference in height between the middle and upper canopy platforms.

| Site | Tree | Middle canopy (m) | Upper canopy (m) | Distance (m) |
| --- | --- | --- | --- | --- |
| Hölstein forest (canopy crane) | 1 | 9.9 | 21.3 | 11.4 |
|  | 2 | 12.7 | 20.7 | 8.0 |
|  | 3 | 12.5 | 19.0 | 6.5 |
| Marburg forest (tree climbers) | 4 | 12.1 | 21.2 | 9.1 |
|  | 5 | 14.9 | 22.9 | 8.0 |
|  | 6 | 13.0 | 21.6 | 8.6 |

#### Canopy crane based canopy placement

The availability of a canopy crane has substantial advantages with regards to getting material and people into the canopy efficiently and proved considerably faster than doing it with tree climbers alone; the canopy placement of the experimental platform was completed in ∼4 hours involving the crane driver and a tree climber. For proper adjustements the tree climber had to leave the gondola to adjust the attachment of the platforms to the tree stems. All material was transported with the crane. The platform was strapped to the stem of the tree with one long side of the platform being flush with the stem. A rubber mat was placed between tree bark and the metal frame of the platform to prevent damage to the tree. The frame was then tightly tied to the stem using two large ratchet straps. A piece of rubber protection was also placed between stem and the metal of the ratchet (see supplementary material). In a second step, two people placed all the pots at ground and in the middle and upper canopy and installed the irrigation system and loggers in 6 hours (see below).

#### Climber based canopy placement

At MOF, the platform placement in the canopy required two tree climbers and one person on the ground. All platform material was pulled up into the trees after setting up a pulley system. The attachment of the platform to the stem of the tree consisted of two ratchet straps with a rubber mat for bark protection and was performed the same way as in Hölstein. However, for an additional measure of safety, since the trees were to be climbed multiple times, the platforms were also held by ropes attached to a rigging plate and a heavy-duty round sling that was tied around the stem above the platform (Figure 1 b, photographs in the supplementary). The platforms were adjusted to be in a level position. Once the platforms were set up (and the irrigation system was in place, see chapter 2.4), plants in 12 L pots were placed on the platforms.Two climbers and one person on the ground were again required to pull the pots into the tree and place them into the empty pots screwed onto the platform. The whole setup took 2 ½ days.

### 2.3 Electricity in the forest

In Hölstein, electricity is available on site, and no further actions had to be taken to power the irrigation system and other required devices (Figure 2a). In Marburg, electricity had to be set up with a solar-powered battery system. We installed one independent system at the base of each tree, which consisted of a waterproof plastic box containing a 120 Ah AGM battery that was connected to a circuit breaker, and a battery control unit with two USB plugs and a solar charge controller. Additionally, the box contained waterproof electric outlets on its outside to power 12V devices, such as the irrigation pump, and it was connected to a solar panel to charge the battery. While the boxes were located at the base of each tree, the solar panels were placed in an adjacent canopy opening receiving more light and were connected to the boxes by long cables (Figure 1b).

**Figure 2.**
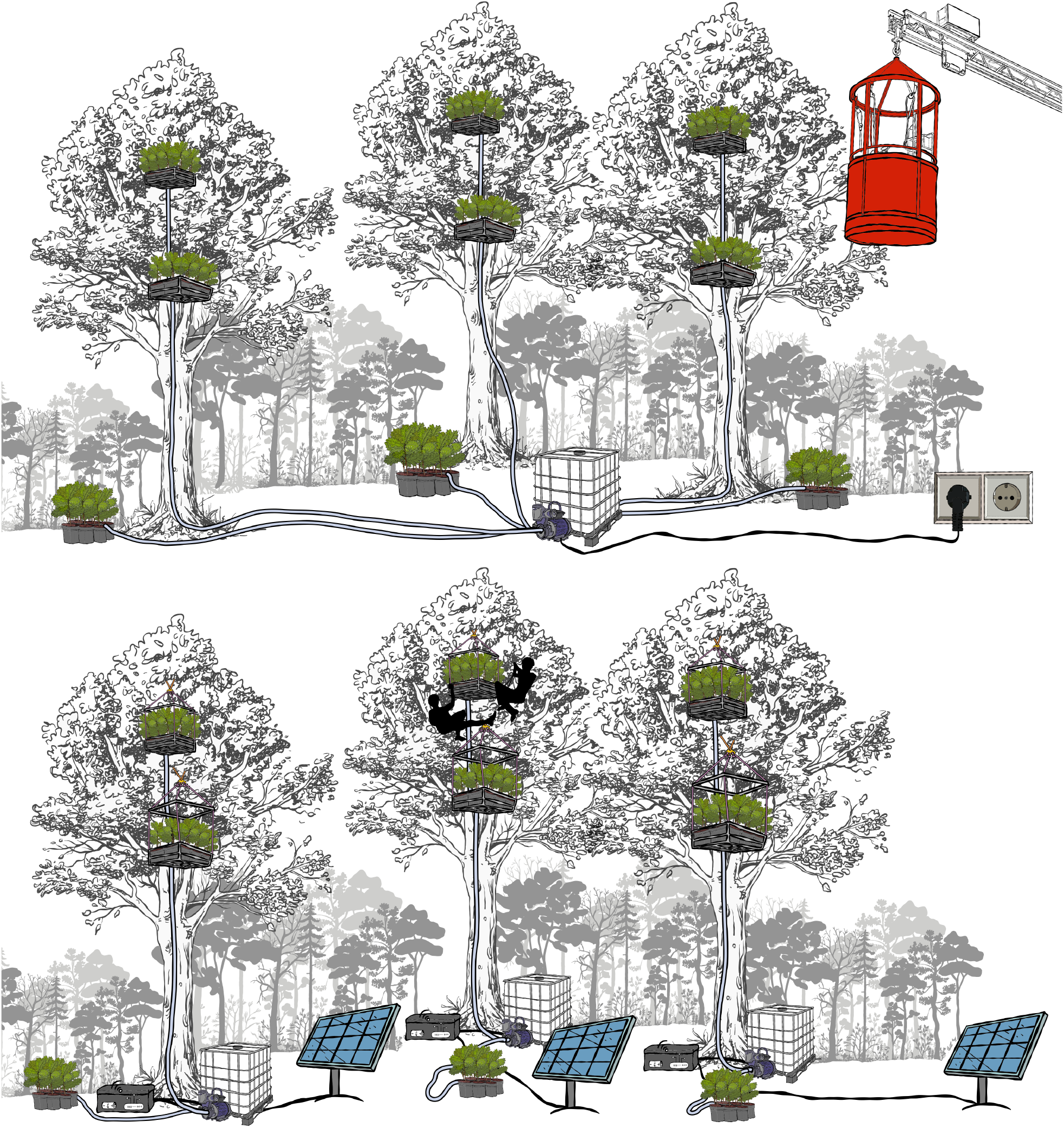
Schematic overview of the TreeTOP experimental platforms installed in the Hölstein Forest (A) and the Marburg Open Forest (B). In both forests, experimental platforms were established at ground, middle, and upper canopy positions to expose tree microcutted-generated saplings to contrasting canopy environments. The Hölstein setup uses a grid-powered irrigation system supplied by a central water reservoir and is delivered by the canopy crane, whereas the Marburg setup relies on battery-powered irrigation systems supplied by individual water reservoirs and maintained by tree climbers.

### 2.4 Canopy irrigation system

Different irrigation systems were used at the two experimental sites based on the resources available within each forest.

#### Irrigation with access to an electric grid

In Hölstein a fully automated irrigation system with a powerful submersible pump and controller including a rain sensor was set up. The irrigation system in Hölstein was custom built by AGRITECH, Hünenberg, Switzerland. One pump with one controller unit (see table S3) was able to water all three trees with their respective platforms, with drippers placed in each pot. They were connected to one central 1000 l bulk container. However, water was not available on site, and the container had to be refilled as needed with the assistance of a local farmer.

#### Battery-based irrigation system

In Marburg, where electricity had to be set up with AGM batteries and solar panels, a less power-consuming solution was needed. An independent watering system was set up at the base of each tree, using a 12 V water pump that could be controlled remotely via a smart switch connected to a mobile router. Since water was not available in the forest, water was filled into black 1000 l bulk containers next to each tree. Black was chosen to prohibit light intrusion and thus algae production. The transport of these containers into the forest was carried out with the help of a tractor, and no refilling was needed throughout the experiment. From the containers, water was pumped through UV-resistant watering hoses with a 19 mm diameter and was then distributed across the six pots of each canopy level using a microdrip system with one dripper per pot.

### 2.5 Microclimatic sensors and irrigtaion control

The same set of waterproof data loggers and sensors was employed at both locations to document microclimatic changes across canopies. Air temperature and light was measured with one logger per canopy level, which was attached to the outside of each platform on the long edge of the frame. The sensors for soil moisture and soil temperature, respectively, were placed within a planting pot, with two pots per canopy height being equipped with the sensors. While air temperature and light were logged on the device and data were read out manually at the end of the experiment, the soil sensors had to be connected to a multi-channel Wi-Fi gateway to transmit their data. In total, three gateways were needed to receive and upload the data from a maximum of eight soil temperature and eight moisture sensors. In MOF, the gateways were placed in the battery boxes at the base of the trees and were connected to a mobile router. In Hölstein, all gateways and the Wi-Fi router were located at the main power station.

### 2.6 Protection from mammals

The Hölstein experimental forest is contained within a fenced area, and no further actions were required to protect material and potted plants on the ground from damage and herbivory by mammals. The MOF is an openly accessible forest. Wire mesh (1.6 m high) was installed around each of the three oak trees to enclose the plants, watering system, and battery boxes.

### 2.7 Statistical analysis

Datasets from all loggers jointly provided information on light intensity, air temperature, and soil temperature in each canopy level. For average daytime measurements, we calculated the mean of measurements between 04:30 in the morning and 21:00 in the evening, while nighttime was defined as the time from 21:00 and 04:30. Differences between canopy levels were determined with a Kruskal-Wallis test and post-hoc Dunn’s test with Benjamini-Hochberg correction.

All statistical analyses were performed in the R environment (version 4.5.1, R Development Core Team, 2021), using the core stats and the rstatix package (Kassambara, 2023). Figures were created using the R-package ggplot2 (Wickham, 2016), ggpubr (Kassambara, 2025), and patchwork (Pederson, 2025). All datasets and accompanying R scripts are available on the PLANTdataHUB (Weil et al., 2023): https://git.nfdi4plants.org/phytoakmeter/treetop_canopy_platform.git.

### 2.8 Budget and parts specifications

Detailed parts lists, quantities and corresponding purchase costs for all components are provided in Tables S2–S6. These include the environmental sensor system (Table S2), the grid-powered irrigation system used in Hölstein (Table S3), the tree-climber platform installation (Table S4), the battery-powered irrigation system used in Marburg (Table S5), and the protective fencing required for the Marburg installation (Table S6). Together, these tables provide a complete overview of the materials required to reproduce both TreeTOP implementations.

The two installations differed primarily in the costs associated with canopy access and irrigation. The Hölstein setup relied on an existing canopy crane, permanent electricity, and a professionally installed grid-powered irrigation system, whereas the Marburg installation used certified tree climbers together with autonomous solar-powered, battery-operated irrigation units. The environmental sensor system was identical at both sites, allowing direct comparison of the monitoring approach despite differences in infrastructure. Canopy access costs (crane operation and tree-climbing personnel) were not included because they depend strongly on local infrastructure and personnel costs.

## 3 Results

### 3.1 Microclimatic conditions across canopy strata

After placing saplings of the *Quercus robur* clone DF159 (Herrmann et al., 1998, Bouffaud et al. 2026) on the platforms at both experimental sites and observing them for one month (from mid-July till mid-August), the sensor data showed changes in microenvironmental conditions across canopy strata in the Marburg and the Hölstein forest. In all trees, air temperature, soil temperature, and light intensities increased with tree height (Figure 3). Mean daytime air temperature was 2.0 °C higher in the upper canopy than on the ground in Hölstein and 2.3 °C higher in Marburg. Likewise, mean daytime light intensity was approximately 12,500 lx higher in Hölstein and 12,200 lx higher in Marburg (Figure 3). In contrast, mean nighttime air temperatures differed by less than 1 °C among canopy strata. No significant difference was observed over mean air temperature per night. For all other measurements, differences between ground and upper canopy were always significant, with additional differences between the ground and middle canopy, as well as between the middle and upper canopy in most cases (Figure 3). Daily temperature extremes differed more strongly among canopy strata than mean temperatures (Figure 4). At both study sites, maximum air temperatures increased consistently from the ground to the upper canopy, resulting in the strongest vertical temperature stratification observed during the study. In contrast, minimum air temperatures differed only slightly among canopy levels, indicating that nocturnal temperatures were comparatively homogeneous throughout the canopy during the sampling period. The same general pattern was observed for soil temperatures, with maximum temperatures increasing more strongly with canopy height than minimum temperatures. However, during night, substrate temperatures were significantly different between canopy levels while air temperatures were non-significant between canopy levels.

**Figure 3.**
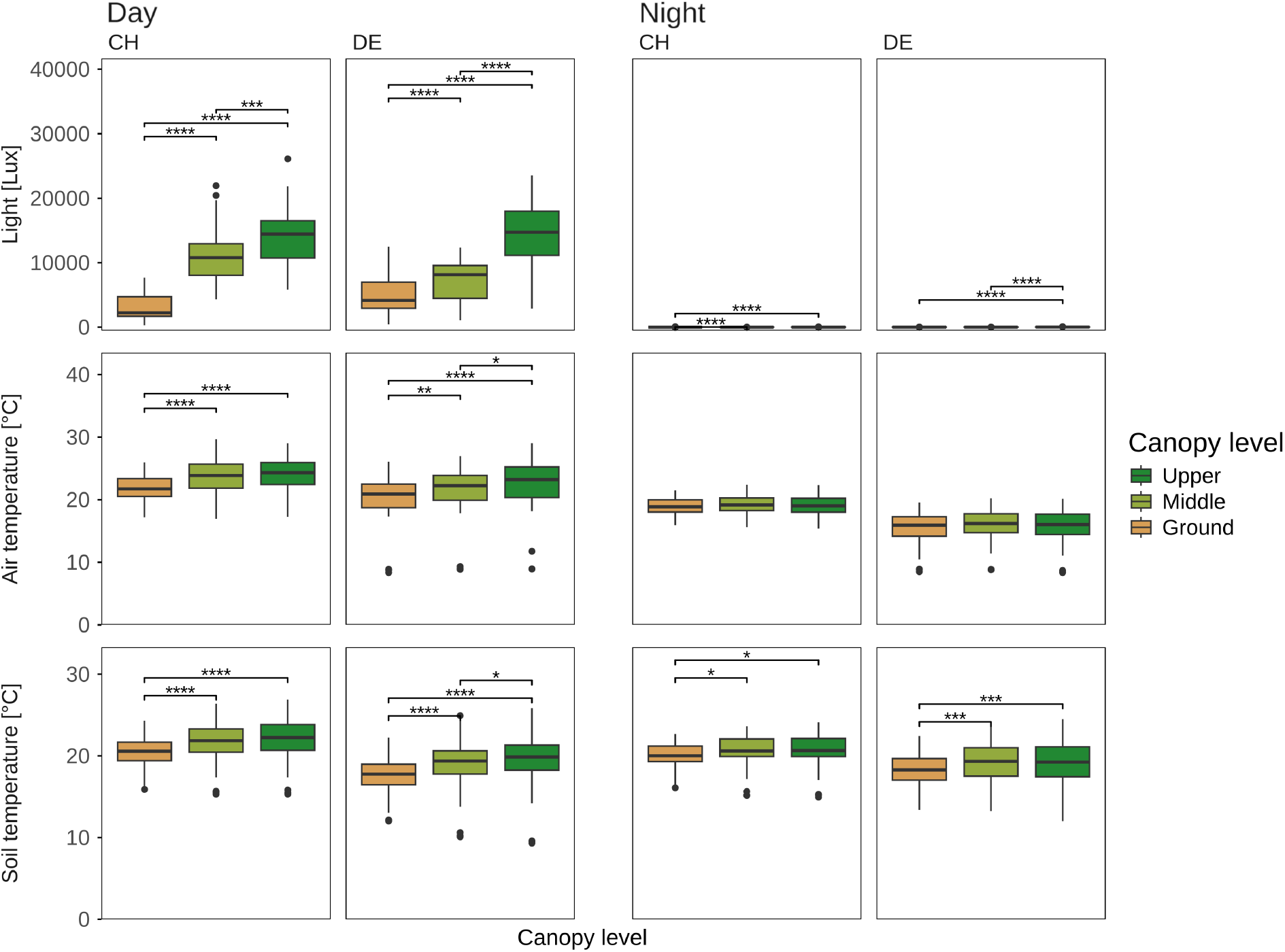
Mean daytime and nighttime light intensity, air temperature, and soil temperature measured at the ground, middle, and upper canopy in the Hölstein Forest (CH, Switzerland) and the Marburg Open Forest (DE, Germany). Differences among canopy levels were assessed using Kruskal–Wallis tests followed by Dunn’s post hoc tests with Benjamini–Hochberg correction for multiple comparisons. Asterisks indicate a significant difference at ^****^*p* < 0.0001 ^***^*p* < 0.001, ^**^*p* < 0.01, ^*\**^p < 0.05

**Figure 4.**
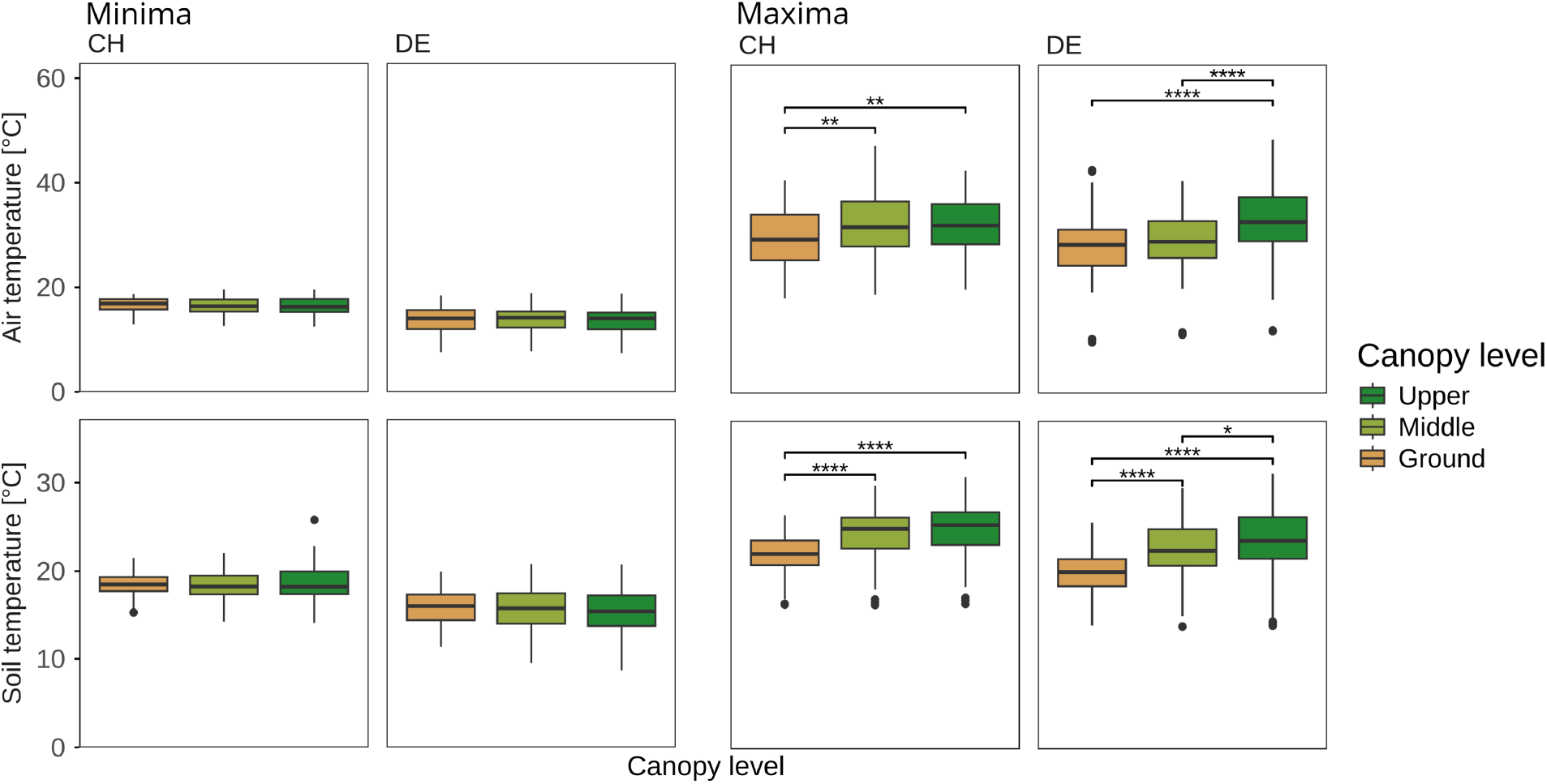
Daily minimum and maximum air and soil temperatures measured at the ground, middle, and upper canopy in the Hölstein Forest (CH, Switzerland) and the Marburg (DE, Marburg) Open Forest. Differences among canopy levels were assessed using Kruskal–Wallis tests followed by Dunn’s post hoc tests with Benjamini–Hochberg correction for multiple comparisons. Significant differences were detected only for daily maximum temperatures. Asterisks indicate significant differences at ^****^P < 0.0001, ^***^P < 0.001, ^**^P < 0.01 and ^*^P < 0.05

Soil moisture patterns differed between the two study sites (Figures S2, S3). In Marburg, substrate moisture generally decreased with canopy height, with consistently wetter pots on the ground than in the middle and upper canopy (Figure S3). In contrast, no consistent vertical pattern was apparent at Hölstein (Figure S2). Unfortunately, intermittent interruptions in soil moisture data transmission limited the temporal overlap among canopy strata, minimizing the robustness of comparisons of soil moisture dynamics in Hölstein.

### 3.2 Diurnal temperature dynamics

Comparing daytime and nighttime measurements revealed that plants at each canopy level experienced a significantly higher air temperature during the day than at night, and the divergence between day and night increased with height (Figure 5). In Hölstein, this difference increased from 2.9 °C on the ground to 5.0 °C in the upper canopy, while in Marburg it increased from 5.1 °C to 7.3 °C. For soil temperature, changes between day and night were not as pronounced as for air temperature, with significant differences in the upper canopy at both locations, the middle canopy in Hölstein, and the ground in Marburg (Figure 5). The average air temperature at night was similar across all canopy heights within locations (Figure 5), but differences became apparent when investigated at a finer temporal resolution. At the beginning of night at 21:00, air temperatures on the ground were significantly lower than in the middle and upper canopies (Figure 5). By the end of night at 04:30, the temperature was similar across all heights. The upper canopy experienced the strongest drop in temperature from start to end of the night, followed by the middle canopy (Table S1). However, this difference in cooling rates was only significant when comparing temperatures on the ground to the middle and upper canopy in Hölstein (Figure 5).

**Figure 5.**
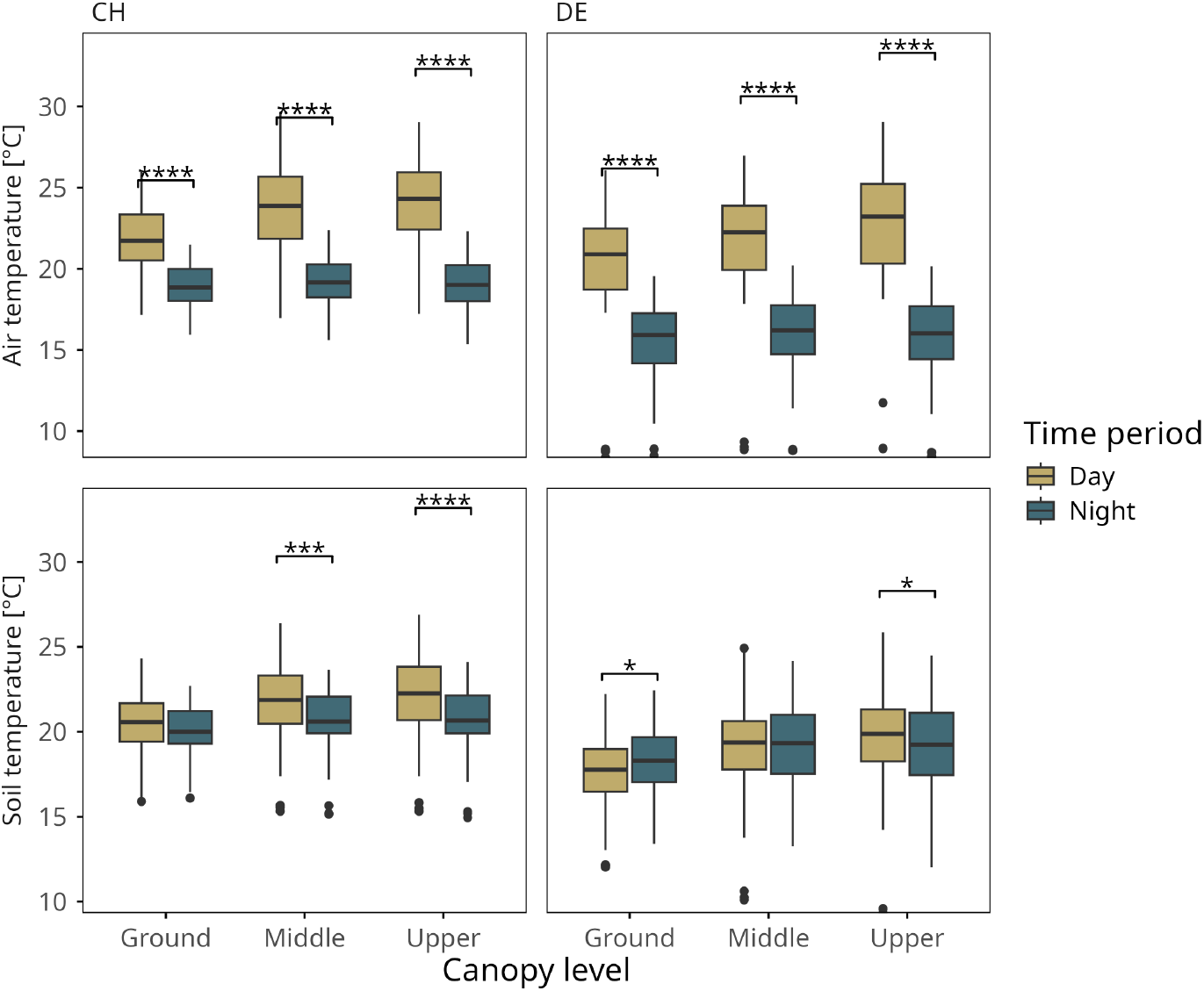
Comparison of mean daytime and nighttime air and soil temperatures measured at the ground, middle, and upper canopy in the Hölstein Forest (CH, Switzerland) and the Marburg Open Forest (DE, Germany). Mean values were calculated separately for each day and night. Differences between daytime and nighttime temperatures within each canopy level were assessed using Kruskal–Wallis tests followed by Dunn’s post hoc tests with Benjamini– Hochberg correction for multiple comparisons. Asterisks indicate a significant difference at ^****^*p* < 0.0001 ^***^*p* < 0.001, ^**^*p* < 0.01, ^*\**^p < 0.05

## 4 Discussion

### 4.1 TreeTOP successfully reproduces canopy microclimatic gradients

TreeTOP successfully reproduced the characteristic vertical microclimatic gradients of mature temperate forest canopies. Light intensity increased strongly with canopy height, while air temperature showed pronounced vertical stratification during the day but much weaker differences at night (Figures 2, 3). These patterns closely resemble those reported from canopy crane studies in other temperate deciduous forests (e.g. Richter et al. 2022; Zahnd et al. 2023) and are consistent with the well-described buffering effect of forest canopies, whereby solar radiation creates strong daytime thermal vertical gradients that largely diminish after sunset (Murakami et al. 2022). Consequently, experimental plants experienced not only realistic average microclimatic differences among canopy strata but also their characteristic diel dynamics.

Particularly noteworthy was that canopy stratification was most pronounced for daily maximum temperatures, whereas daily minima differed little among canopy strata (Figure 4). This indicates that vertical temperature differences were primarily generated during periods of intense solar radiation, while nocturnal cooling largely equalized temperatures across canopy levels. This pattern is consistent with the buffering capacity of forest canopies, which dampens temperature extremes and reduces thermal contrasts during the night (Murakami et al. 2022). From an experimental perspective, this is highly desirable because many physiological responses of plants and their associated organisms are driven by short-term exposure to extreme temperatures rather than by average conditions alone (De Frenne et al. 2024; Teskey et al. 2015). TreeTOP therefore reproduces not only differences in mean canopy climate but also the thermal extremes that are expected to shape canopy processes under climate change.

An inherent limitation of TreeTOP is that the root environment is transferred into the canopy together with the experimental plants. Consequently, both mean and maximum substrate temperatures increased with canopy height (Figures 3, 4), whereas the root systems of mature trees remain in comparatively buffered forest soils. During platform development, we evaluated several approaches to reduce substrate heating, including reflective pot covers and passive water cooling. However, pre-trials showed that neither approach substantially reduced substrate temperatures while remaining practical for long-term field deployment.

Substrate moisture, however, is less clear-cut. In mature trees, foliage in the upper canopy experiences higher radiation, vapor pressure deficit, and transpiration, while water must simultaneously be transported over longer hydraulic pathways with greater hydraulic resistance (Ryan et al. 2006). Consequently, upper-canopy foliage commonly operates under greater hydraulic constraints than leaves lower in the crown. In TreeTOP, lower substrate moisture in upper-canopy pots (Figures S2, S3) may therefore partly reflect these biologically realistic differences in plant water relations rather than representing solely an experimental artefact, although irrigation strategy also contributed to the observed patterns. Whether substrate moisture should be maintained at similar levels across canopy strata or allowed to diverge ultimately depends on the experimental objective. Studies aiming to isolate the effects of atmospheric microclimate should standardize irrigation accordingly, whereas experiments investigating whole-plant responses to canopy environments may intentionally permit differences in substrate moisture to develop.

### 4.2 TreeTOP across contrasting research infrastructures

The successful implementation of TreeTOP in two research forests with contrasting infrastructures demonstrates that the platform is not tied to a particular type of canopy access or technical environment. Rather, the two installations illustrate complementary strategies that can be adapted to the available infrastructure, logistical constraints and research objectives. At Hölstein, the availability of a canopy crane and a permanent electricity supply enabled rapid installation and a centrally controlled irrigation system. Grid-powered irrigation minimized maintenance requirements and allowed automated watering schedules to be implemented throughout the experiment. Such infrastructures are particularly advantageous for long-term experiments, studies requiring frequent access to the canopy, or installations involving large numbers of experimental units. In contrast, the Marburg Open Forest demonstrates that comparable experiments can also be established in forests lacking permanent canopy access or electrical infrastructure. Although installation and maintenance required certified tree climbers and battery-powered irrigation systems, these constraints were largely offset by the greater flexibility in platform placement. Unlike canopy cranes, which are limited by their operational radius and canopy accessibility, climbing allows platforms to be installed in virtually any suitable tree, greatly expanding the range of forests in which TreeTOP can be deployed. Because only a small number of research forests worldwide provide permanent canopy cranes or comparable infrastructure, the ability to establish autonomous experimental platforms using climbing techniques substantially broadens the applicability of manipulative canopy experiments. Researchers can therefore select the installation strategy according to local infrastructure without fundamentally changing the experimental design, facilitating comparable studies across forest types and geographic regions.

### 4.3 New opportunities for mechanistic canopy ecology

TreeTOP enables a range of manipulative experiments that have previously been difficult to implement in mature forest canopies. Recent syntheses have highlighted the need for a mechanistic understanding of how canopy microclimates mediate forest responses to global change and influence ecological processes across forest ecosystems (De Frenne et al. 2021; Verheyen et al. 2024). Standardized plants can be established across canopy strata to investigate how microclimate influences plant performance, phenology, species interactions and the assembly of tree-associated microbiomes under realistic field conditions. Experimental manipulations such as altered irrigation, nutrient addition, shading or warming can be applied independently or in combination, while reciprocal transplant experiments among canopy strata or between forest sites provide a powerful approach for disentangling the relative importance of microclimate, host identity and dispersal in shaping canopy-associated communities (Arnold et al. 2025; Saueressig et al. 2026).

The modular design of TreeTOP also facilitates the integration of modern sensor technologies. Time-lapse cameras and automated imaging systems, as currently developed within projects such as PhytOakmeter, could continuously quantify plant growth, phenology and canopy development throughout the growing season. Future developments may further enable automated monitoring of herbivory, providing continuous observations of plant–herbivore interactions under contrasting microclimatic conditions. Combined with environmental sensors measuring light, temperature, soil moisture and additional variables such as vapor pressure deficit, these systems would enable real-time links between canopy microclimate and plant physiological responses.

Perhaps the greatest strength of TreeTOP is that it transforms mature forest canopies from observational systems into experimentally tractable environments. Rather than describing naturally occurring patterns alone, researchers can manipulate environmental conditions, continuously monitor organismal responses and directly test ecological mechanisms across realistic canopy gradients. Because the same experimental design can be implemented in research forests with or without permanent canopy infrastructure, TreeTOP also provides a framework for coordinated multi-site experiments across forest types, climates and geographic regions.

### 4.4 Limitations and lessons learned for experimental canopy ecology

Although TreeTOP proved robust under field conditions, the pilot implementation identified several practical limitations that should be considered when establishing similar experiments. In climber-based installations, maintenance visits should be kept to a minimum. At the Marburg Open Forest, repeated climbing activities occasionally damaged irrigation hoses, resulting in water leakage and additional maintenance climbs. Most of these problems could likely be avoided by installing all components within a tree during a single climbing session and limiting subsequent canopy access to essential inspections.

Reliable irrigation is essential for long-term canopy experiments. The two irrigation concepts performed comparably well and maintained healthy experimental plants throughout the study. However, the pilot implementation in the Marburg Open Forest identified two aspects that should be improved in future battery-powered installations. First, only a single non-return valve was installed between the ground and middle canopy platforms, allowing water to drain from the upper canopy into the middle canopy after irrigation and reducing the amount of water delivered to the upper platform. This issue was avoided in the Hölstein installation by placing non-return valves between each canopy level. Second, manual decisions on irrigation frequency introduced unnecessary variation in substrate moisture. Future deployments should therefore standardize irrigation schedules across sites and, where possible, automate watering based on predefined schedules or sensor feedback. The solar-powered battery system in Marburg nevertheless provided sufficient energy for irrigation, data transmission and remote monitoring throughout the experiment. Experiments requiring substantially higher irrigation frequencies or prolonged operation under low solar radiation, however, may require larger battery capacities or replacement batteries.

The sensor network proved sufficiently robust for long-term monitoring but highlighted several opportunities for improvement. Remote monitoring substantially reduced maintenance effort and enabled rapid detection of technical failures, including the replacement of a malfunctioning Wi-Fi switch in Marburg. At Hölstein, communication between the Ecowitt gateways and Wi-Fi router was generally reliable, but intermittent transmission failures occurred for the soil moisture sensors, whereas soil temperature data were received consistently. Although the underlying cause remains unclear, future deployments spanning larger distances between trees may benefit from additional gateways, more powerful routers or redundant communication pathways. The choice of environmental sensors should reflect the objectives and available infrastructure of the experiment. We deliberately selected a low-cost, low-maintenance monitoring system in which air temperature and light data were stored on HOBO loggers and retrieved manually after the experiment. Although this minimized installation effort, maintenance and power consumption, sites with permanent electricity and internet access may benefit from fully automated sensor networks capable of continuously transmitting additional variables such as wind speed, precipitation or vapor pressure deficit.

Finally, platform orientation should be standardized wherever possible. In practice, however, the exact position of each platform was constrained by crown architecture and, at Hölstein, by crane accessibility. Although all platforms were preferentially installed on the north-western side of the stem, some variation in orientation was unavoidable and should be considered when comparing experiments across trees or sites. Overall, the practical limitations identified during this pilot implementation can readily be addressed through minor modifications of the experimental design and should not limit the broader applicability of TreeTOP.

## 5 Conclusions

TreeTOP provides a standardized and transferable platform for experimental canopy ecology. The successful implementation in two research forests with contrasting infrastructures demonstrates that realistic manipulative experiments can be established across mature forest canopies without permanent canopy infrastructure. By combining standardized experimental units with naturally occurring canopy microclimatic gradients, TreeTOP creates new opportunities to investigate plant performance, species interactions and microbiome assembly under realistic field conditions. We expect that the platform will facilitate coordinated experiments across forest types and climatic regions, advancing mechanistic understanding of canopy ecology in a changing world.

## Author contributions

The PhytOakmeter consortium: design of the experiment, review of the manuscript, aquisition of funding. Julia Baumeister: Writing of the manuscript, planning and implementation of TreeTOP in Marburg forest, data curation, analysis, and visualisation. Moe Bakhtiari: Review of the manuscript, planning and implementation of TreeTOP in Basel forest, data curation. Mona Schreiber: Review of the manuscript, planning and implementation of TreeTOP in Marburg forest, visualisation of experimental setup. Michael Eisenring: Review of the manuscript, planning of TreeTOP in Basel forest, funding acquisition. Martin Gossner: Review of the manuscript, planning of TreeTOP in Basel forest, funding acquisition. Susanne Walden: planning of TreeTOP in Marburg forest. Sylvie Herrmann: Review of the manuscript, conceptualisation of the study. Francois Buscot: Review of the manuscript, conceptualisation of the study. Katrin Heer: Review of the manuscript, conceptualisation of the study, funding aquisition. Lars Opgenoorth: Writing of the manuscript, conceptualisation of the study, funding acquisition. All authors reviewed the manuscript.

## Acknowledgements

This work was carried out within the framework of PhytOakmeter, funded by the Deutsche Forschungsgemeinschaft (DFG, German Research Foundation) – 507084794 and the Swiss National Science Foundation (SNSF) - 310030E_215899 / 1. Work in Switzerland was conducted at the Swiss Canopy Crane II research site of the University of Basel in Hölstein. We thank Ansgar Kahmen and Günter Hoch for accommodating our experiment at the Hölstein research site. We thank Dennis Handte for assisting with the crane operation and the installation of the experimental platforms in Hölstein, and Anouchka Perret-Gentil for assistance in the field. Work in Germany was carried out within the Marburg Open Forest (MOF), a research forest of Philipps-University in Marburg. We thank Moe for tree climbing, planning and logistics support. We ensured data FAIRness through the use of the DataPLANT infrastructure, including the personal assistance network and the integrated tool stack, with the DataHUB serving as the central component.

