## Supplementary material for "TreeTOP: Plant experimental platforms in canopy space"

### Supplementary Information

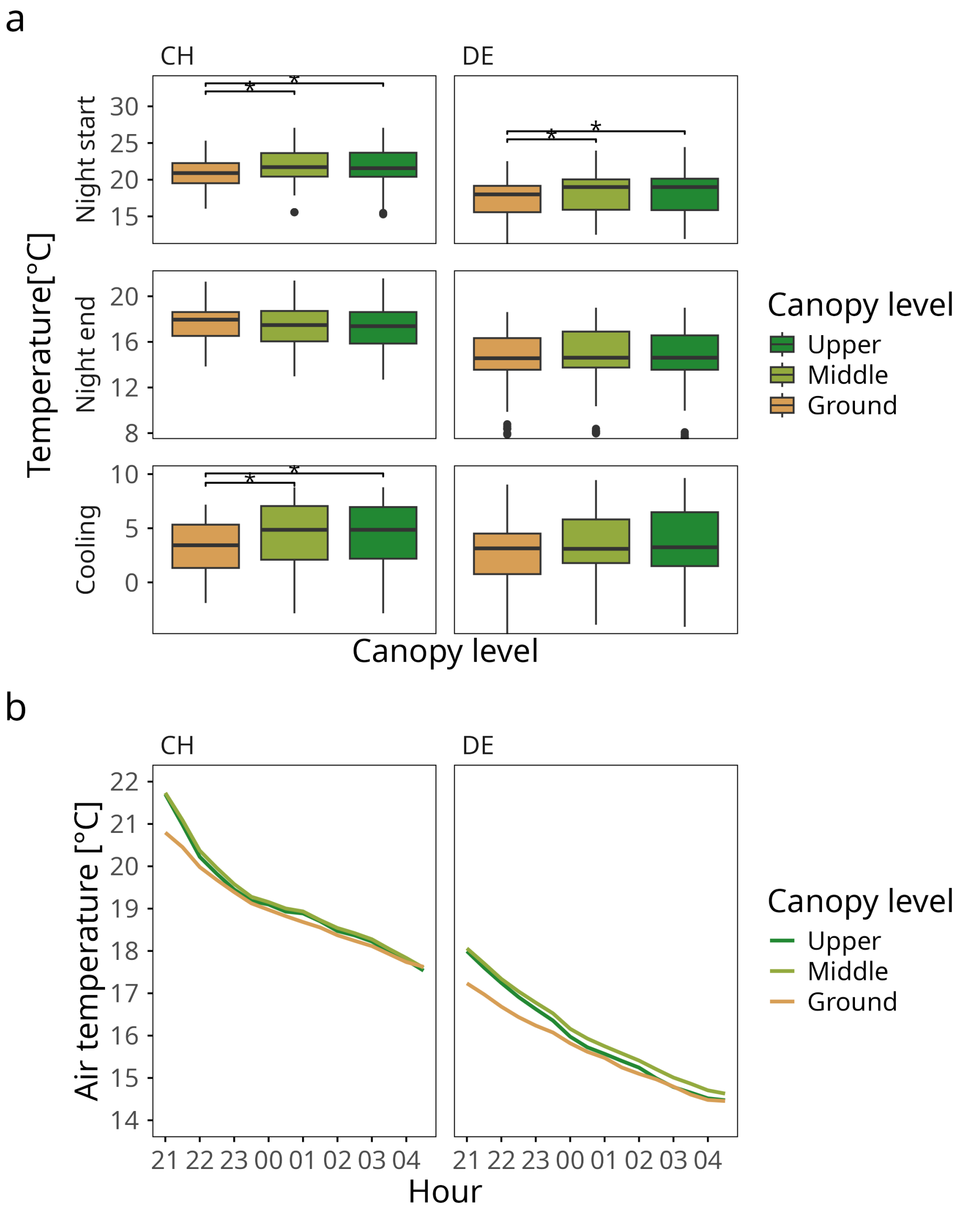


**Figure S2:** Changes of air temperature during the night a) Temperature variation recorded across canopy levels at the start and the end of each night. The start of nighttime was defined as 21:00, while the end of nighttime was set to 04:30. The cooling temperature shows the difference between temperatures measured at the start and end. b) Average air temperature during the night in 30-minute intervals measured on the ground, the middle canopy and the upper canopy at the two TreeTOP locations in the Hölstein forest (CH, Switzerland) and the Marburg Open Forest (DE, Germany), respectively. Differences in a) are shown after Kruskal-Wallis test and post-hoc Dunn's test with Benjamini-Hochberg corrected *p* values. Asterisks indicate a significant difference at $^*p < 0.05$

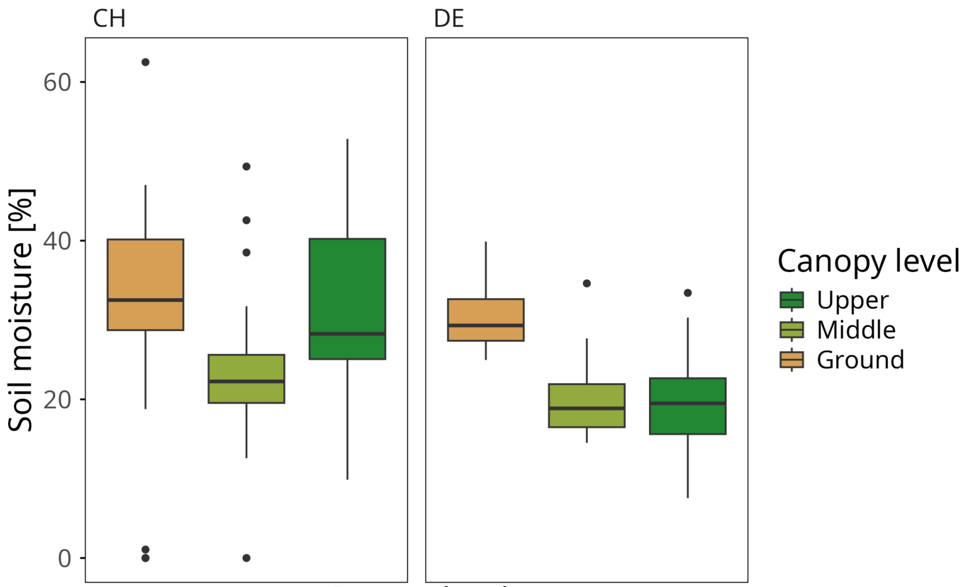


**Figure S3.** Distribution of daily mean soil moisture measured at ground, middle, and upper canopy TreeTOP platforms in the Hölstein Forest (CH, Switzerland) and the Marburg Open Forest (DE, Germany). Boxes show the median, interquartile range, and 1.5 × interquartile range, with points indicating outliers.

A B

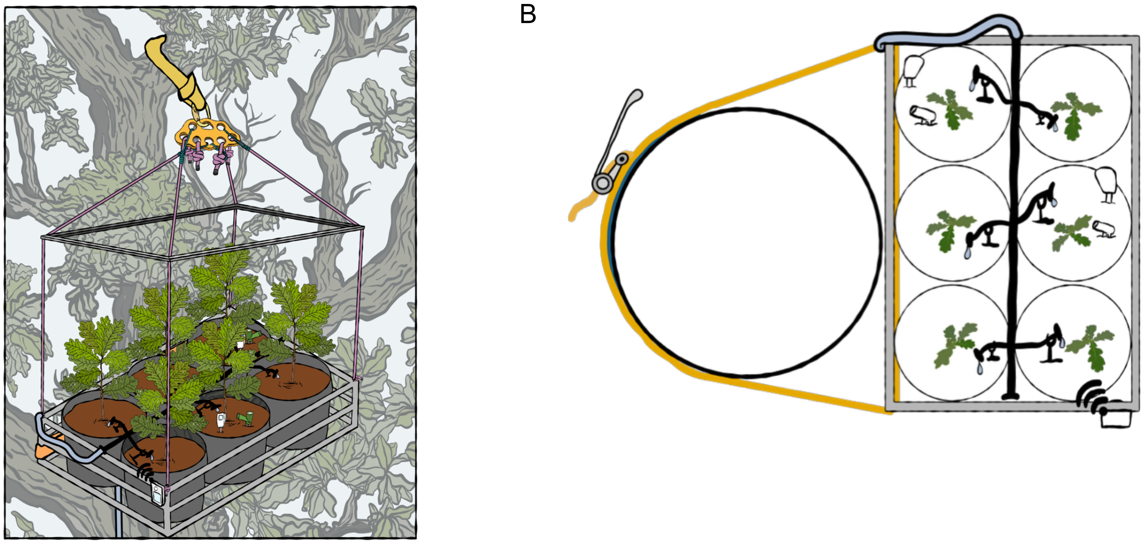

**Figure S4:** Schematic overview of the TreeTOP platform. (A) Three-dimensional view of the experimental platform suspended from an overhead branch using a heavy-duty round sling, rigging plate, ropes, and locking carabiners, while being secured to the tree stem with ratchet straps and protective rubber mats. The platform accommodates six experimental pots equipped with drip irrigation, soil temperature and moisture sensors, and air temperature/light loggers mounted on the platform frame. (B) Top view showing the attachment of the platform to the tree stem, the arrangement of the six pots, the main irrigation hose with pressure-compensated drippers, and the positions of the environmental sensors.

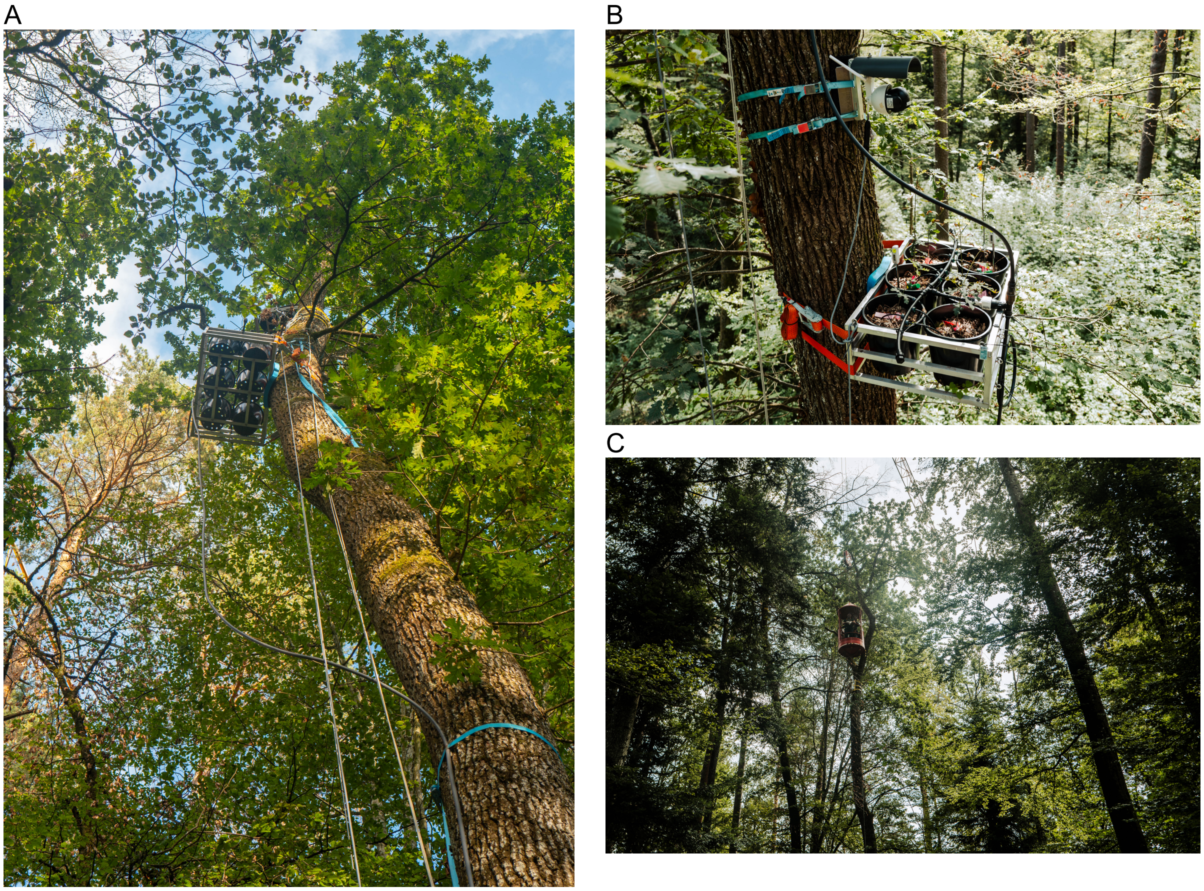

**Figure S5.** TreeTOP Middle Platform installed in the Hölstein research forest (Switzerland) using the Swiss Canopy Crane II. (A) TreeTOP platform mounted in the upper canopy and secured to the tree stem with ratchet straps, while suspended from an overhead branch using tree-rigging equipment. Irrigation tubing supplies water from a ground-based reservoir. (B) Close-up of the experimental platform showing the six planted pots, drip irrigation system, environmental sensors, and data loggers attached to the platform frame. (C) Canopy crane gondola (Swiss Canopy Crane II) providing access to the experimental platforms during installation and maintenance. Photos by M. Eisenring.

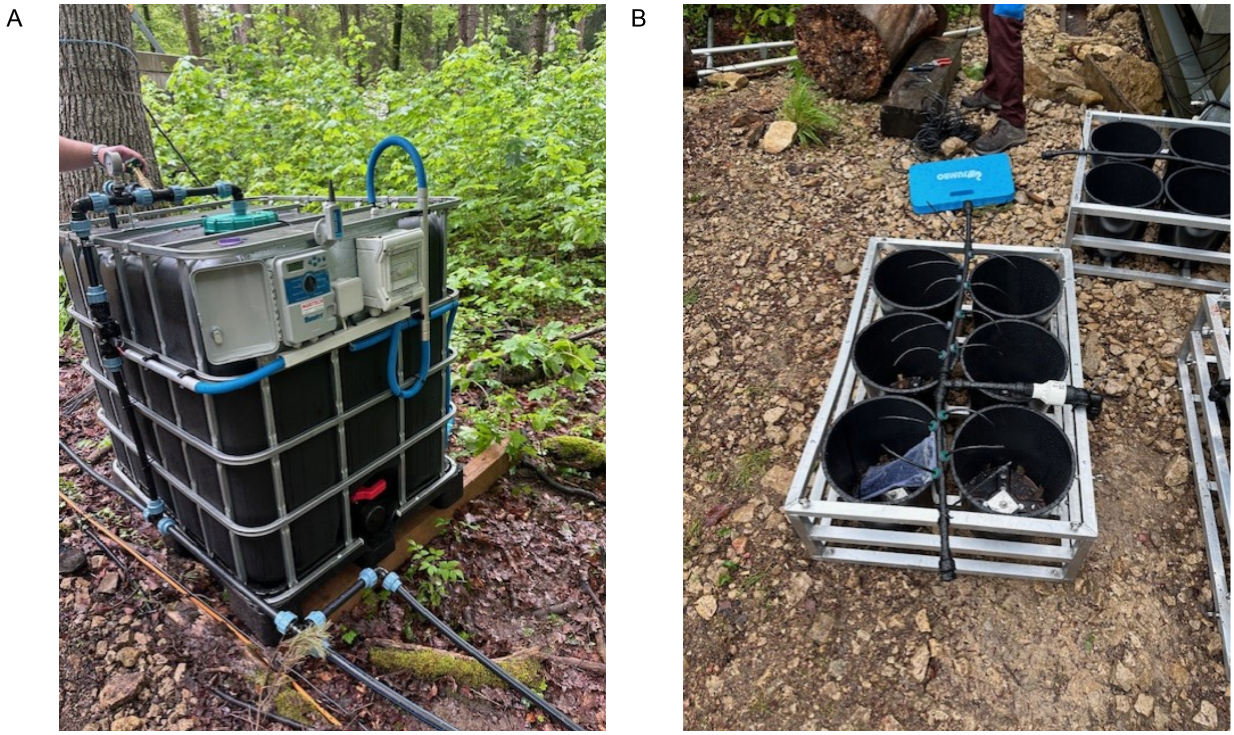


**Figure S6.** Grid-powered irrigation system installed in the Hölstein Forest. (A) 1000 L intermediate bulk container (IBC) equipped with a submersible pump, pump controller, and irrigation controller supplying water to the TreeTOP platforms. (B) Example of a TreeTOP platform showing the main irrigation line, connecting pipe, pressure-compensating drippers, and soil moisture sensor installed before deployment into the canopy.

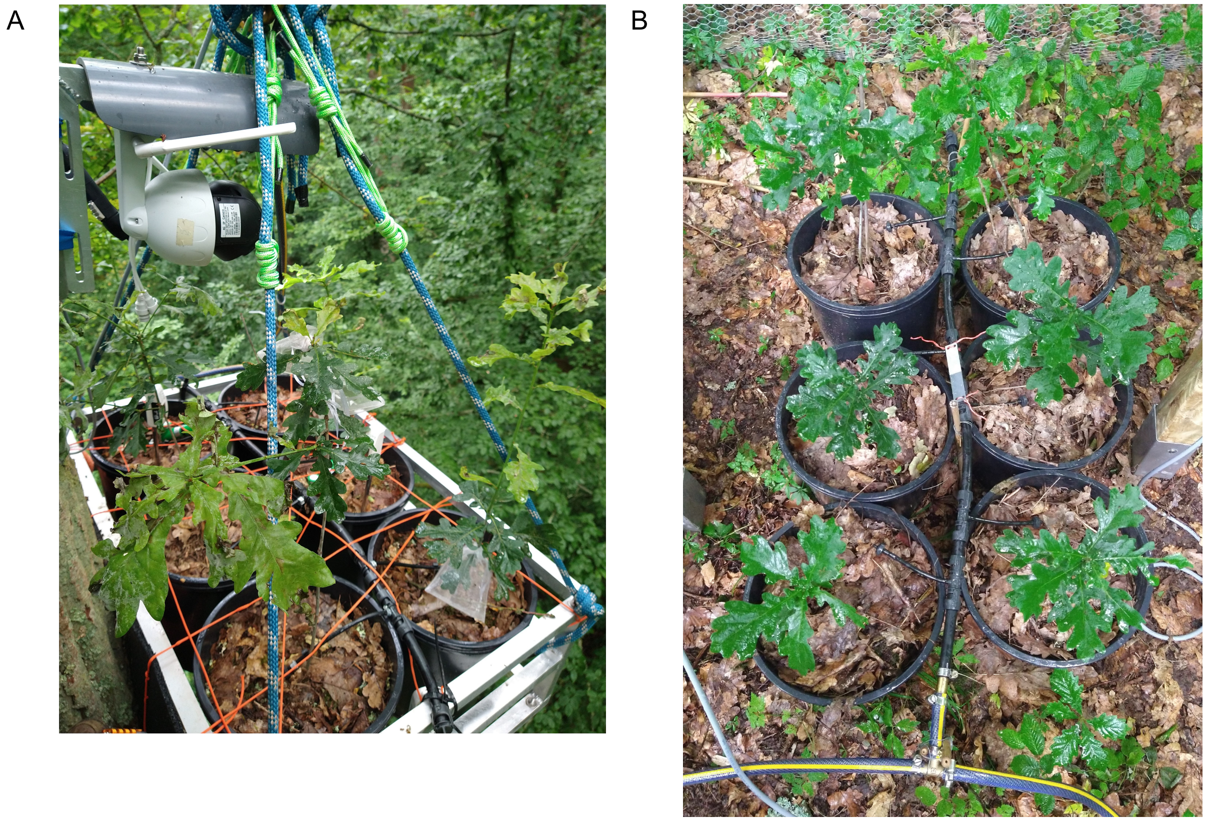


**Figure S7.** TreeTOP platforms installed in the Marburg Open Forest . (A) TreeTOP platform installed in the middle canopy, equipped with six experimental plants, drip irrigation, and environmental sensors. (B) Ground-level TreeTOP platform showing the arrangement of the six experimental plants and the drip irrigation system prior to deployment.

**Table S1:** Mean air temperature at the beginning (21:00) and end (04:30) of the night across canopy strata in Hölstein Forest (Switzerland) and Marburg Open Forest (Germany). Cooling was calculated as the mean temperature decrease between the beginning and end of each night.

| Site | Canopy level | Air temperature at start of night [°C] | Air temperature at end of night [°C] | Cooling [°C] |
|---|---|---|---|---|
| **Hölstein Forest** | Ground | 20.87 | 17.62 | 03.28 |
| | Middle | 21.89 | 17.61 | 04.31 |
| | Upper | 21.88 | 17.54 | 04.37 |
| **Marburg Open Forest** | Ground | 17.29 | 14.46 | 02.66 |
| | Middle | 18.14 | 14.63 | 03.19 |
| | Upper | 18.08 | 14.47 | 03.28 |

**Table S2.** Components required for the environmental sensor system installed at both Hölstein Forest (Switzerland) and Marburg Open Forest (Germany). The table lists the required number of components per site, their function within the monitoring system, and the corresponding purchase costs.

| Product | No./site | No. total | Purpose | Price [€] |
|---|---|---|---|---|
| **HOBO Pendant Temperature/Light 64K Data Logger** | 9 | 18 | Monitoring of air temperature and light on each platform | 93 |
| **HOBO BASE-U-4 universal optic USB base station** | 1 | 2 | Read out of data from the HOBO loggers | 175 |
| **Ecowitt wireless soil moisture sensor WH51** | 18 | 36 | Monitoring of soil moisture in two pots per platform | 15 |
| **Ecowitt wireless soil temperature sensor WN34S** | 18 | 36 | Monitoring of soil temperature in two pots per platform | 15 |
| **Ecowitt Wi-Fi gateway GW1200** | 4 | 8 | Receiving data from the soil sensors and uploading them to a server | 28 |
| **Mobile router 4G Wi-Fi hotspot** | 1 | 2 | Providing internet for the Ecowitt gateways and the watering system | 38 |
| **SIM card for mobile data** | 1 | 2 | Inserted into the mobile router, 5 GB mobile data per month | 7 |

**Table S3.** Components required for the installation of the grid-powered irrigation system supporting nine TreeTOP platforms in the Hölstein Forest. The system in Hölstein was custom built by AGRITECH, Hünenberg, Switzerland. The table lists the required components, their function within the irrigation system, and the corresponding purchase or installation costs. All costs are rounded to the nearest full Euro.

| Product | No. | Purpose | Price [€] |
|---|---|---|---|
| **Hunter 4 station X-core irrigation computer** | 1 | Irrigation controller | 775 |
| **Hunter X-CORE residential Irrigation water Pump control system** | 1 | Material needed to control the pump | 1628 |
| **Hunter X-CORE residential Irrigation Submersible pump** | 1 | Pump with unidirectional valve placed inside the water container | 1447 |
| **25 mm water pipe** | 150 m | Main water pipes from the container to the trees | 635 |
| **16 mm water pipe** | 100 m | Thinner pipes installed | 238 |
| **Drip irrigation per platform (custom made)** | 9 | 12 microdrippers, 1 non-return valve per tree level | 720 |
| **Setup costs** | 1 | Planning and installation by an external company | 3583 |
| **1000 L black intermediate bulk container (IBC)** | 1 | Water storage | 636 |
| **NAAN Click Tif dripper green 8 L/h** | 20 | Drip irrigation | 12 |
| **Spiess ANTELCO "Asta Clip Stake" 14 cm** | 25 | Holding the drippers in place inside the pots | 13 |

**Table S4:** Components required for the tree-climber-based installation of six TreeTOP platforms in the Marburg Open Forest . The table lists the required components, their function within the platform suspension system, and the corresponding purchase costs. All costs are rounded to the nearest full Euro.

| Product | No. | Purpose | Price [€] |
|---|---|---|---|
| **Ratchet straps 6 m x 55 mm** | 12 | Attaching platforms to the tree stem | 46 |
| **Petzl PAW rigging plate** | 6 | Pulling the platform up into the tree and attaching it to the round sling | 54 |
| **Heavy-duty round sling, 1.5-3 m usable length** | 6 | Carrying the weight of the platform together with the ratchet straps, chosen length depends on branch thickness and platform location | 25 - 41 |
| **Steel carabiner or steel shackle** | 6 | Connecting piece between rigging plate and round sling. The shackle needs to be secured with wire to prevent an accidental opening, the carabiner needs to have a locking mechanism | 24 or 4 |
| **Rubber mat, 230 cm x 115 cm x 8 mm, cut to fit** | | Protective layer between ratchet or platform and tree bark | 30 |

**Table S5.** Components required for the installation of the battery-powered irrigation system supporting nine TreeTOP platforms in the Marburg Open Forest. The table lists the required components, their function within the irrigation system, and the corresponding purchase costs. All costs are rounded to the nearest full Euro.

| Product | No. | Purpose | Price [€] |
|---|---|---|---|
| **Shurflo 12 V pump, 3 bar, model 2088-403-144** | 3 | Watering of all platforms per tree | 131 |
| **Shelly Plus 1 Wi-Fi switching actuator 16 A** | 3 | Activation and deactivation of the pump | 17 |
| **1000 L black intermediate bulk container (IBC)** | 3 | Water storage | 140 |
| **Gardena microdrip system (list all components?)** | 9 | Watering of pots | |
| **12 V AGM batteries, 120 Ah** | 3 | Electricity for the pump and Ecowitt gateways | 148 |
| **Non-return valve** | 3 | Preventing water backflow from the middle canopy to the ground | 15 |
| **LUX-TOOLS PRO watering hose 13mm, 50m** | 3 | Watering pipe going from the IBC to platforms on the ground, middle and upper canopy of each tree | 60 |
| **10 pc Gardena endline drippers** | 6 | Drip irrigation, pressure-compensating, 2 L/h | 9 |
| **Gardena End Plug 13 mm, 5pc** | 2 | Plug attached to the end of the connecting pipe on each platform | 5 |
| **Gardena connecting pipe flex 13 mm, 15m** | 1 (70 cm per platform) | Pipe connecting all 6 pots per platform | 14 |
| **Gardena drip irrigation pipe 4.6mm, 15m** | 1 (100 cm per platform) | Drip irrigation line per pot, attached to the connecting pipe | 9 |
| **Pipe Pegs 4.6 mm, 10pc** | 6 | Holding the drippers in place inside the pots | 7 |
| **Gardena T-joint connectors 13mm (1/2") – 4.6 mm (3/16"), 5pc** | 11 | Connectors between connection pipe and drip irrigation pipe | 8 |
| **Hose coupling 13 mm** | 9 | Connecting watering hose with the drip irrigation connecting pipe | 4 |
| **13 mm T-joints** | 9 | Watering hose connections at each platform intersection | |
| **Hose clamps 10-16 mm** | | | |

**Table S6.** Components required for constructing protective fencing around the TreeTOP platforms in the Marburg Open Forest. The table lists the required components, their function, and the corresponding purchase costs. All costs are rounded to the nearest full Euro.

| Product | No. | Purpose | Price [€] |
|---|---|---|---|
| **Wire mesh** | | Protection from damage | |
| **Wooden post 180 cm x 7 cm x 7 cm** | 18 | Holding the wire mesh in place | 11 |
| **Impact ground sleeve 750 mm x 71mmx 71mm** | 18 | Post anchor | 5 |
